# Non-Viral Engineering of CAR-NK and CAR-T cells using the *Tc Buster* Transposon System™

**DOI:** 10.1101/2021.08.02.454772

**Authors:** Emily J. Pomeroy, Walker S. Lahr, Jae Woong Chang, Joshua Krueger, Bryce J. Wick, Nicholas J. Slipek, Joseph G. Skeate, Beau R. Webber, Branden S. Moriarity

**Affiliations:** Department of Pediatrics, University of Minnesota; Masonic Cancer Center, University of Minnesota; Center for Genome Engineering, University of Minnesota; Stem Cell Institute, University of Minnesota

## Abstract

Cancer immunotherapy using T cells and NK cells modified with viral vectors to express a chimeric antigen receptor (CAR) has shown remarkable efficacy in treating hematological malignancies in clinical trials. However, viral vectors are limited in their cargo size capacity, and large-scale manufacturing for clinical use remains complex and cost prohibitive. As an alternative, CAR delivery via DNA transposon engineering is a superior and cost-effective production method. Engineering via transposition is accomplished using a two-component system: a plasmid containing a gene expression cassette flanked by transposon inverted terminal repeats (ITRs) paired with a transposase enzyme that binds to the ITRs, excises the transposon from the plasmid, and stably integrates the transposon into the genome.

Here, we used the newly developed hyperactive *Tc Buster* (Bio-Techne) transposon system to deliver a transposon containing a multicistronic expression cassette (CD19-CAR, mutant DHFR, and EGFP) to primary human peripheral blood (PB) NK cells and T cells. We optimized methods to avoid DNA toxicity and maximize efficiency. Our cargo contained a mutant dihydrofolate reductase (DHFR) which allowed us to enrich for stable transposon integration using methotrexate (MTX) selection. We then tested CAR-NK and CAR-T cells in functional assays against CD19-expressing Raji cells. CAR-expressing NK and T cells produced significantly more cytokines than CAR-negative controls and efficiently killed target cells. We recognize that cryopreservation manufactured CAR-expressing cells will be necessary for clinical translation. We observed reduced cytotoxicity of CAR-NK cells immediately after thaw, but increasing the NK dose overcame this loss of function.

Our work provides a platform for robust delivery of multicistronic, large cargo via transposition to primary human NK and T cells. We demonstrate that CAR-expressing cells can be enriched using MTX selection, while maintaining high viability and function. This non-viral approach represents a versatile, safe, and cost-effective option for the manufacture of CAR-NK and CAR-T cells compared to viral delivery.

## Introduction

DNA transposons are natural DNA transfer vehicles that can be used for DNA delivery into human cells. In nature, they exist as well-defined elements in which the transposase gene is flanked by inverted terminal repeats (ITRs) that encode transposase binding sites. They can be used as a tool for stable genomic insertion by flanking an expression cassette with ITRs and co-delivering the transposase enzyme via an expression plasmid, mRNA, or protein^1^.

Several DNA transposons have been used in such a manner in mammalian cells. In the 1990s, the *Sleeping Beauty* (*SB*) transposon system was molecularly reconstructed by eliminating inactivating mutations found in members of the Tc1/mariner family of transposons isolated from fish^2^. This reactivated transposon system has since been used for stable gene transfer and insertional mutagenesis in many vertebrate cell types, including human cells. Subsequently, the *piggyBac* (*PB*) and *Tol2* transposable elements were isolated from insects and fish, respectively, and have been optimized for enhanced activity in mammalian cells^3,4^. *SB, PB*, and *Tol2* can all be used as efficient non-viral tools for stable gene delivery, and each of these has been used for gene delivery in primary human lymphocytes^5^. The hAT superfamily of transposons (including *Tol2*) are found in diverse species^6^. A novel representative of this family is *Tc Buster*, originally isolated from the red flour beetle^7^. *Tc Buster* has been shown to be active in human cell lines and has a comparable transposition efficiency to *SB* and *PB*^6,8^. A hyperactive mutant of *Tc Buster* was recently developed (Bio-Techne, Minneapolis, MN) for the non-viral manufacture of cellular therapies.

Transposons have many meaningful advantages as an alternative to viral vectors for stable gene transfer. Several clinical gene therapy products have been developed using CD34+ hematopoietic stem cells or T cells genetically modified using recombinant viruses; namely γ-retroviruses, and lentiviruses^**9–19**^. These delivery methods carry the risk of insertional mutagenesis via activation of proto-oncogenes or inactivation of tumor suppressor genes^**20–22**^. In addition, large-scale manufacturing of these viral vectors for clinical use is cost-prohibitive, inconsistent, and impedes progression through clinical trials; particularly for academic institutions and emerging biotech companies. The use of transposon systems has been pursued as an alternative to viral vectors due to rapid, reproducible, and cost-effective production and a favorable safety profile^**10,20,23**^. However, any vector that integrates into chromosomes poses the risk of insertional mutagenesis. A comparative study of the target site integration properties of *SB* and *PB* transposons as well as gammaretroviral and lentiviral systems in primary human CD4+ T cells ranked their safety profiles based on multiple criteria, including distance from the 5’-end of any gene and distance from any cancer-related gene. This analysis established *SB* as having the most favorable integration profile, followed by *PB*, suggesting that engineering via transposition is a safer alternative to engineering with viral vectors^**24**^. Further, genome wide analysis comparing integrations of *Tc Buster* to *piggyBac* and *Sleeping Beauty* revealed a unique but comparable integration pattern for *Tc Buster*^6^.

The use of transposon systems for gene delivery in human lymphocytes has been most widely studied as a method for generating human T cells engineered to express CARs^**25,26**^. Clinically, the *SB* system has been used to introduce CD19-specific CARs to patient- and donor-derived T cells^**27,28**^. Many preclinical studies and clinical trials thus far have introduced *SB* transposase and CAR by electroporation of bulk peripheral blood mononuclear cells (PBMCs)^**29–33**^. CAR-expressing T cells were subsequently expanded over several weeks in culture using feeder cells engineered to express the target antigen and co-stimulatory molecules^**27**^. Efforts are now being made to shorten the culture time before patient infusion (NCT04102436, NCT04289220). One such trial is underway in which PBMCs are transferred into the patient within 2 days after electroporation with *SB* transposase, CD19-CAR, and membrane-bound IL15 (NCT03579888). Signaling through the CAR and mbIL15 gives genetically-modified T cells a selective advantage after transplant, fostering their outgrowth^**34**^.

Another approach for shortening culture time *ex vivo* is to electroporate T cells directly, rather than as bulk PBMCs from which T cells need to be selected. This has been challenging as the delivery of transposons via plasmids leads to DNA toxicity in T cells due to a type I interferon (IFN) response^**35,36**^. Thus, efforts have been made to minimize the amount of DNA introduced to the T cell. Minicircle vectors are DNA plasmid delivery vehicles that do not carry a bacterial origin of replication or bacterial resistance genes, reducing the size of the vector to only that of the expression cassette^**37**^. This approach has been used to achieve stable expression of transgenes in primary human T cells with efficiencies over 50 percent^**35,38,39**^. However, minicircle vectors rely on a complex and inefficient recombination-based removal of the plasmid bacterial regions prior to plasmid purification, often leading to low plasmid yield, bacterial genomic DNA contamination, and overall inconsistency between productions. More recently, Nanoplasmids (Nature Technology) have been developed that contain a small (<500 bp) backbone that encodes the origin of replication from the Rep/iteron plasmid R6K^**40**^ and the RNA-OUT (antisense RNA) selectable marker^**41**^. This minimal backbone reduces DNA toxicity and thus nanoplasmids can be made with high yield and without complex purification methods^**41,42**^.

Thus far, the use of transposons for NK cells has been mostly applied to the NK-92 cell line^**43**^. Recently, the *SB* transposon system has been used to deliver a CAR to cytokine-induced killer cells for targeting CD33 on chemoresistance acute myeloid leukemia (AML) in patient-derived xenografts^**1,44**^. However, lessons can be learned from T cells on the use of transposons for CAR delivery to primary NK cells. The initial approach of electroporating PBMCs with transposon-based CARs suggests that NK cells could be selectively outgrown instead of T cells^**29,30**^. Indeed, some reports have shown outgrowth of NK cells reaching 50% of the PBMC population after co-culture with feeder cells^**30**^. Thus, this approach could be optimized for selection of CAR-expressing NK cells, or delivery of a mixed population of CAR-T and -NK cells might be advantageous as NK cells have been shown to produce inflammatory cytokines to help shape the adaptive immune response^**45**^.

Alternatively, the use of minicircle or nanoplasmid vectors to deliver transposons directly to purified NK cells is an attractive option. NK cells share many properties with T cells, and delivery of DNA to NK cells has been shown to induce similar toxicity^**46**^. Thus, reducing the amount of DNA delivery by using mRNA-encoded transposase in combination with a minicircle- or nanoplasmid-encoded transposon may be ideal.

The use of transposons for engineering NK cells is not limited to the delivery of CARs. Other modifications have been proposed to enhance aspects of NK cell activity including persistence, migration, and cytotoxicity. This includes the introduction of self-stimulating cytokine receptors^**47**^, strong activating receptors, or dominant negative versions of NK cell inhibitory receptors^**48**^. Such modifications could be used in combination with CAR delivery to create an NK cell expertly equipped to kill a broad range of tumor types. Transposons provide a non-viral strategy to introduce all of these transgenes in one vector that could be scaled up for clinical use. Here we developed such a method for clinically scalable production of CAR-NK and CAR-T cells using the *Tc Buster* Transposon System. We achieve manufacturing of enriched (>99%+; 46%+ pre-MTX selection) CAR-expressing cells in a matter of 3 weeks and show functional efficacy of CD19 targeting CAR-NK and CAR-T cells. Our work provides a versatile, cost-effective, and clinically scalable platform for the manufacture of engineered NK and T cells.

## Results

### Optimized Engineering of CAR-NK Cells using Hyperactive Tc Buster

In an effort develop a robust platform for the manufacture and enrichment of CAR-expressing NK and T cells, we used the newly developed hyperactive *Tc Buster*™ (TcB-M, Bio-Techne) system to deliver a nanoplasmid containing a CD19-CAR-DHFR-EGFP expression cassette (3.7 kb transposon, Figure 1A) to primary human peripheral blood (PB) NK cells.

**Figure 1.**
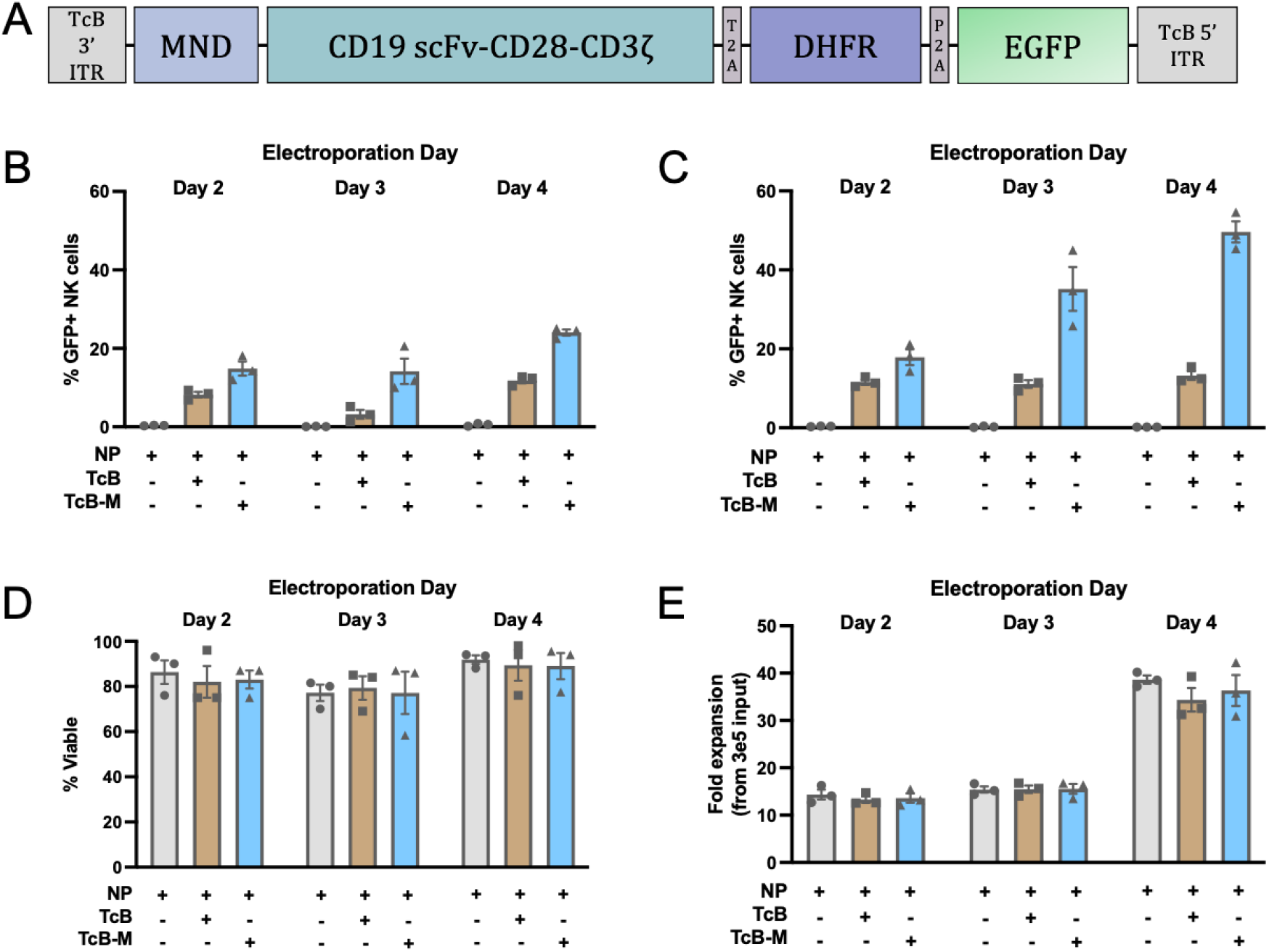
Delivery of a CD19-CAR-DHFR-EGFP transposon to NK cells using the *Tc Buster* Transposon System. (A) The 3.7 kb transposon flanked by *Tc Buster* ITRs, containing an MND promoter, second generation CD19 CAR, methotrexate resistant DHFR mutant, and enhanced GFP. Elements are separated by 2A ribosomal skip sequences. This cargo was cloned into a nanoplasmid (NP) backbone for delivery. (B-E) Primary human peripheral blood (PB) NK cells (n=3 human donors) were expanded for 2, 3, or 4 days with mbIL21- and 41BBL-expressing K562 feeder cells at a 2:1 (feeder:NK) ratio. NK cells were electroporated with the nanoplasmid transposon (NP) alone or in combination with mRNA encoding either *Tc Buster* (TcB) or the hyperactive mutant *Tc Buster* (TcB-M). Two days after electroporation, NK cells were expanded with feeder cells (5:1 feeder:NK ratio) for 1 week to allow for the loss of transient NP expression. After expansion, GFP expression was measured by flow cytometry, viability was measured by Trypan Blue exclusion, and fold expansion was calculated. (B) NK cells electroporated without pre-treatment with RNase inhibitor. (C-E) NK cells electroporated after 5-minute pre-treatment with RNase inhibitor.

We previously developed a method for efficient RNA delivery of CRISPR-Cas9 to primary human peripheral blood (PB) NK cells^**49**^. We used this as a starting point for optimization of transposition. Briefly, we stimulated NK cells by co-culture with membrane bound IL21 (mbIL21)- and 41BBL-expressing K562 feeder cells for 7 days before electroporating them with mRNA encoding Cas9 and chemically modified sgRNA using the Neon Transfection System^**49**^. We initially followed this same process to deliver the nanoplasmid transposon and mRNA encoding *Tc Buster* transposase. We observed low delivery efficiency as measured by GFP expression (2.07 ± 0.37%) and poor recovery (12.00 ± 0.58%) of electroporated cells (Supplemental Figure 1A and 1B).

Suspecting that cytosolic DNA sensors may be upregulated in NK cells during the activation process leading to poor delivery efficiency and cell recovery^**50**^, we surmised that delivering DNA earlier in the activation process may precede the upregulation of DNA sensors. Thus, we tested delivery of the transposon with *Tc Buster* (TcB) or the hyperactive *Tc Buster* mutant transposase (TcB-M) on day 2, 3, or 4 of feeder cell-mediated activation (Figure 1B). In line with our hypothesis, we observed higher transposition efficiency at earlier time points, with the highest efficiency on day 4 of activation (11.85 ± 1.26% for TcB and 24.05 ± 1.36% for TcB-M). Importantly, the majority of feeder cells in the population are eliminated by day 4 of the activation process (Supplemental Figure 2).

Activation of NK cells is known to lead to the upregulation of ribonucleases^**51**^, and as we deliver the transposase as mRNA, we tested the treatment of NK cells with a ribonuclease inhibitor (RNase A, RNase B, and RNase T2) for 5 minutes prior to electroporation with transposase mRNA (Figure 1C – 1E). We found that the addition of the RNase inhibitor enhanced transposition efficiency, and again observed the highest transposition efficiency when electroporation was performed on day 4 of NK cell activation (Figure 1C, 13.23 ± 1.89% for TcB and 49.63 ± 4.64% for TcB-M). We re-expanded the NK cells 48-hours after electroporation for one week and compared viability and fold-expansion for each electroporation time-point. While differences in cell viability were minimal (Figure 1D), NK cells electroporated on day 4 of activation had significantly higher fold-expansion than those electroporated on day 2 or day 3 (Figure 1E). Thus, optimal stable transposition in NK cells was achieved by electroporation on day 4 of expansion and pre-treatment with RNase inhibitor before electroporation.

### Enrichment of CAR-NK Cells using Methotrexate

The antifolate methotrexate (MTX) inhibits wild-type dihydrofolate reductase (DHFR), which is essential for cell growth and proliferation^**52,53**^. Our cargo contained a MTX-resistant DHFR mutant (L22F, F31S)^**54**^, allowing us to select and expand NK cells that had undergone stable transposon integration. This approach is being used clinically to enrich for CAR-expressing T cells (NCT04483778). We incorporated this selection step into our production process during an additional round of feeder cell-mediated expansion (Figure 2A). Three doses of MTX were evaluated to determine the minimal dose needed to kill control cells and enrich engineered cells while maintaining high cell viability and recovery (Figure 2B and 2C). An optimal dose of 250 nM MTX completely killed control cells (Figure 2B) and enriched cells engineered with either TcB or TcB-M to >99% GFP+ (Figure 2C). CD19-CAR expression was confirmed by staining cells with Atto 647N-labeled recombinant human CD19 (Supplemental Figure 3A). Our optimized protocol achieves manufacturing in ∼20 days and results in 99.2% (± 0.5%) CAR+ NK cells expanded 1380-fold (± 104.4) from electroporation input (Figure 2D).

**Figure 2.**
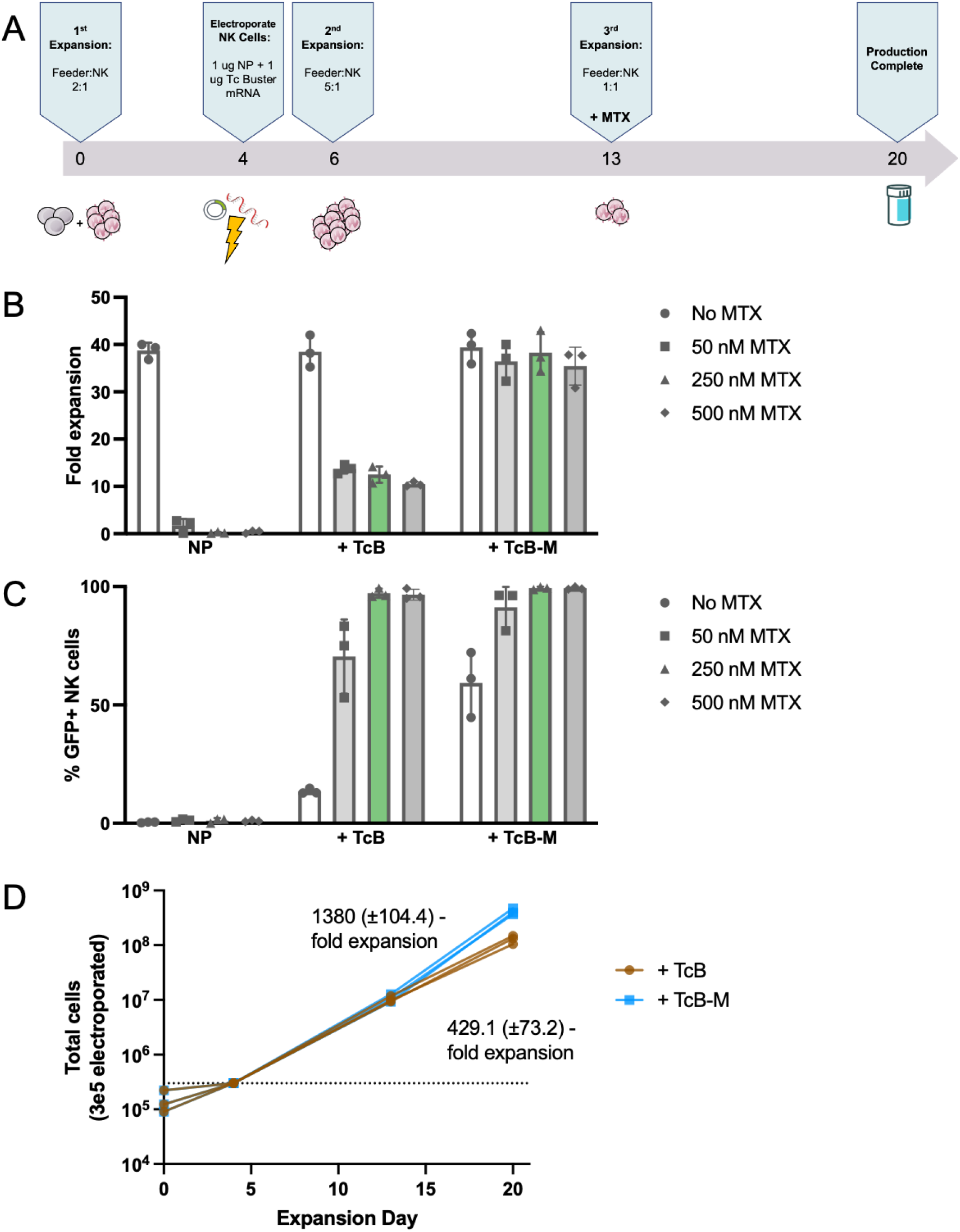
Efficient production of CAR-NK cells using an optimized engineering pipeline incorporating MTX selection. (A) Timeline for production of CAR-NK cells: Primary human peripheral blood NK cells (n=3 human donors) are activated with mbIL21- and 41BBL-expressing K562 feeder cells at a 2:1 (feeder:NK) ratio for 4 days. On day 4, NK cells are treated with RNase inhibitors and electroporated with transposition reagents. Two days after electroporation, NK cells are expanded for one week using a 5:1 (feeder:NK) ratio. NK cells are expanded again for one week at a 1:1 (feeder:NK) ratio in media containing MTX. (B) NK cells were expanded a third time in media containing increasing doses of MTX and fold expansion was calculated from input into the third expansion in order to determine optimal MTX selection dose. (C) GFP expression was measured by flow cytometry. (D) Total cells were tracked over the course of the full production timeline and fold expansion was calculated from electroporation input.

### Functional Validation of CAR-NK Cells

We evaluated NK cells engineered with TcB-M and selected to >99% CAR+ with MTX for testing in functional assays against the CD19-expressing Raji Burkitt’s lymphoma cell line. NK cells electroporated with the transposon nanoplasmid alone (without transposase) and expanded without MTX selection served as CAR-negative controls. CAR-negative or CAR-positive NK cells were co-cultured with Raji target cells for 5 hours at various effector-to-target (E:T) ratios and analyzed for expression of markers of activation and cytotoxicity. CAR-NK cells produced more inflammatory cytokines IFNγ and TNFα than CAR-negative NK cells after co-culture (Figure 3A and 3B). CAR-NK cells also expressed more CD107a on their surface, a marker of degranulation (Figure 3C). In a luciferase-based killing assay, CAR-NK cells killed over 90% of Raji cells in 24 hours at the lowest E:T ratio of 1:3 (Figure 3D). These data show robust functionality of CAR-NK cells against CD19-expressing target cells.

**Figure 3.**
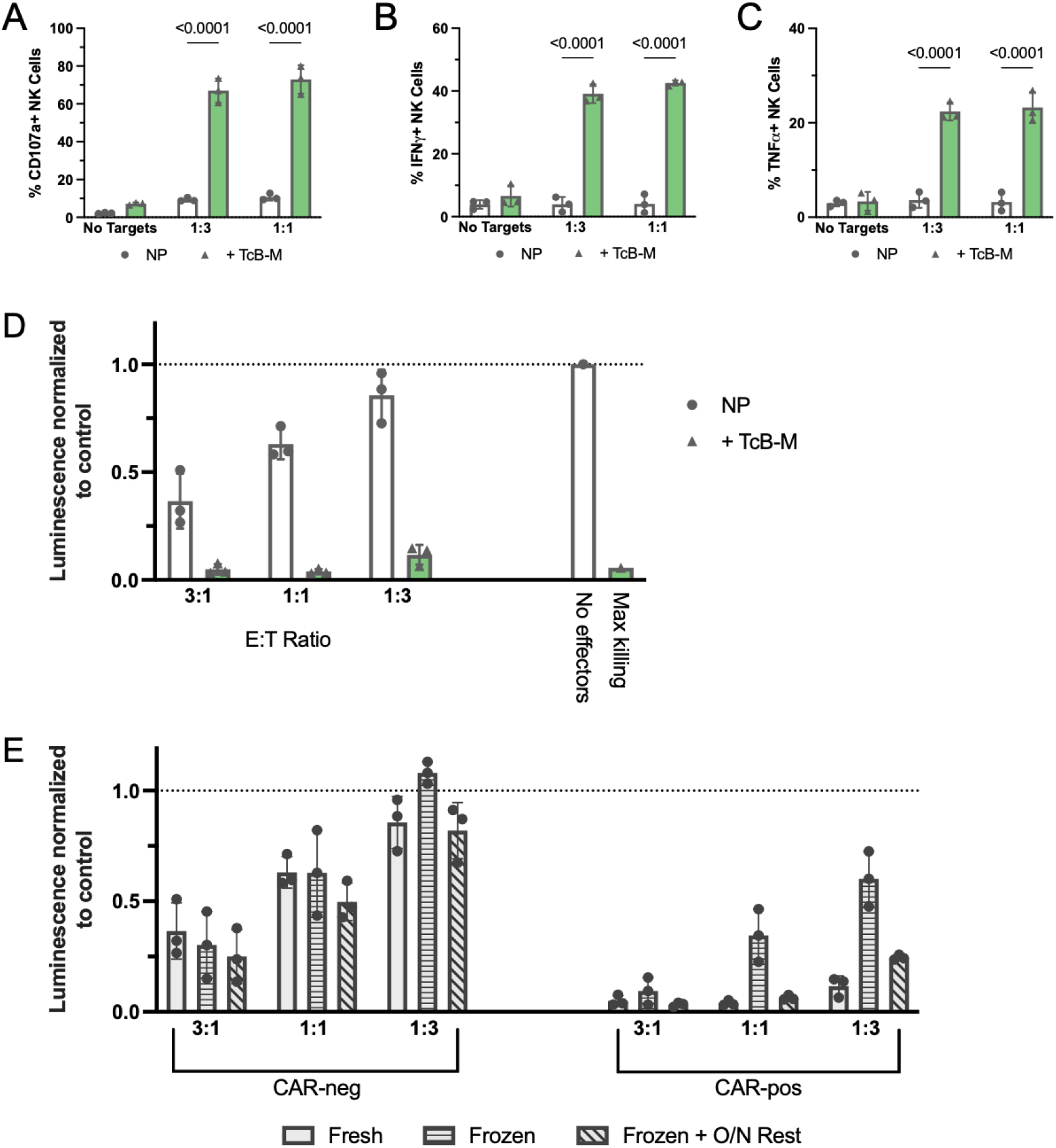
CD19-CAR-expressing NK cells show enhanced activity and tumor cell killing compared to CAR-negative controls. CAR-positive cells were engineered with TcB-M and selected with 250 nM MTX. CAR-negative control cells were electroporated with NP alone and were not selected with MTX. (A-C) CAR-positive and CAR-negative NK cells were co-cultured with CD19+ Raji cells at the indicated effector-to-target (E:T) ratios for 5 hours. Co-cultures were set up in triplicate. After co-culture, NK cells were analyzed by intracellular flow cytometry for the expression of CD107a (A), IFNγ (B), and TNFα (C). (D-E) CAR-positive and CAR-negative NK cells were co-cultured with luciferase-expressing CD19+ Raji cells at the indicated E:T ratios for 24 hours. Co-cultures were set up in quadruplicate. After co-culture, cells were incubated with d-luciferin for 10 minutes and luminescence was read on a plate reader. The assay was performed with fresh NK cells immediately after production and with cryopreserved NK cells immediately after thaw or after an overnight rest in media containing 100 IU/mL IL2.

Our ultimate goal is to deploy this manufacturing pipeline for clinical use, requiring cryopreservation and banking of manufactured CAR-NK cells. Recent studies have shown that while cryopreservation may have little effect on NK viability, it can cause a loss of cytotoxic function^**55**^. Thus, we tested cytotoxicity of CAR-NK cells immediately after thaw or after overnight culture in media containing 100 IU/mL IL2 (Figure 3D). Although cytotoxicity of CAR-NK cells immediately after thaw was reduced compared to fresh cells, target cell killing was still efficient at higher E:T ratios (3:1). Cytotoxicity of frozen CAR-NK cells was restored to that of fresh cells after an overnight rest in media containing IL2. We did not observe a reduction in cell number after the overnight rest, but we did observe a slight reduction in cell viability (Supplemental Figure 4A and 4B). These data provide further evidence for loss of NK function from cryopreservation, but suggest that allowing NK cells to recover after thaw or delivering higher doses of NK cells are approaches to circumvent this loss of function.

### Efficient Engineering of CAR-T Cells using Tc Buster Transposition

A long-term goal of our work is to develop a combination therapy of CAR-NK and CAR-T cells. Thus, we next evaluated the *Tc Buster* transposon system for the manufacture of CAR-T cells. We stimulated CD3+ primary human T cells with αCD3/αCD28 DynaBeads for 2 days and electroporated them with the nanoplasmid alone or in combination with TcB or TcB-M mRNA. Three days post-electroporation, we selected for transposon integration with 250 nM MTX for an additional 7 days, for a total production timeline of 12 days (Figure 4A). We observed successful enrichment (>99% CAR+) of engineered cells with MTX while maintaining high cell viability (Figure 4B – 4D). We similarly tested CAR-T cells in a killing assay against CD19-expressing Raji cells and observed near complete killing of target cells at higher E:T ratios (3:1) (Figure 4E).

**Figure 4.**
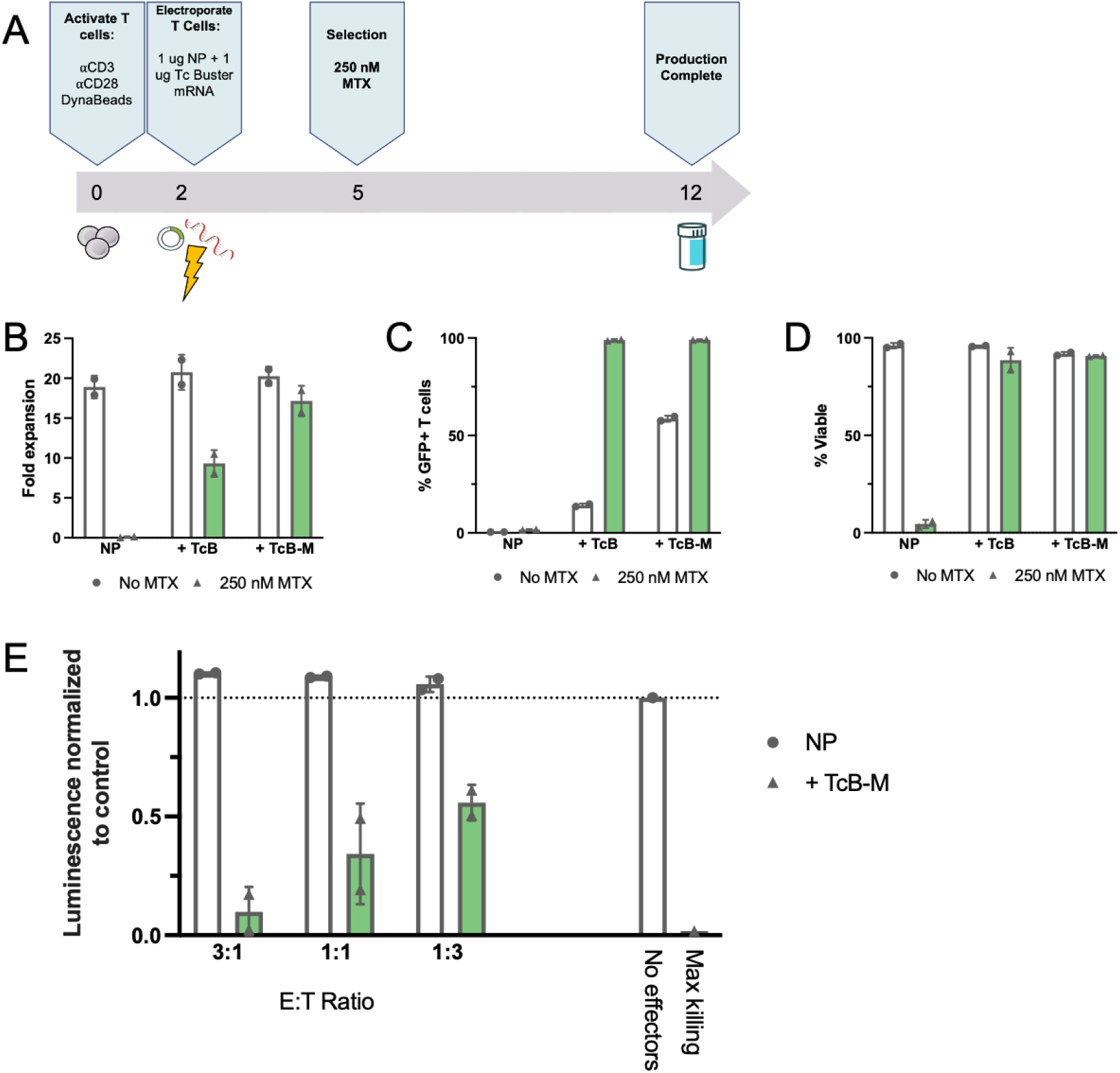
CD19-CAR T cells can be generated using *Tc Buster* transposition and have functional killing against tumor targets. (A) Timeline for production of CAR-T cells: Primary human peripheral blood T cells are activated with αCD3/αCD28 DynaBeads for 2 days. On day 2, DynaBeads are removed, and T cells are electroporated with transposition reagents and subsequently returned to DynaBead-containing cultures. 3 days after electroporation, MTX is added to the media at a concentration of 250 nM. 7 days after MTX selection, production is complete and T cells are cryopreserved. (B-D) Production was performed with and without MTX selection. At the end of production, T cells were counted and fold expansion from electroporation input was calculated (B), GFP expression was measured by flow cytometry (C), and viability was measured by trypan blue exclusion (D). CAR-positive cells were engineered with TcB-M and selected with 250 nM MTX. CAR-negative control cells were electroporated with NP alone and were not selected with MTX. CAR-positive and CAR-negative T cells were co-cultured with luciferase-expressing CD19+ Raji cells at the indicated E:T ratios for 24 hours. Co-cultures were set up in quadruplicate (n=2 donors). Co-cultures were incubated with D-luciferin for 10 minutes and luminescence was read on a plate reader.

## Discussion

Current methods for the manufacture of CAR-NK and CAR-T cells for clinical use largely rely on viral transduction. The nature of NK cells as first responders to viral infection has likely led them to evolve resistance to viral infection and explains their intransigence to viral transduction^**56**^. In addition, the use of viral vectors for CAR-NK and CAR-T production has several downsides. Production and handling of viral vectors is time consuming and costly, and viral vectors are limited in the size and complexity of their cargo and carry the risk of insertional mutagenesis. Engineering using transposons is an attractive alternative approach as production is convenient and cost-effective, and transposons have been shown to have a safer integration profile with reduced preference for integration near gene regulatory elements^**24**^.

The goal of our work was to develop a protocol for manufacturing CAR-NK and CAR-T cells using the *Tc Buster* transposon system. The *Tc Buster* transposon is a member of the hAT superfamily of transposons and was originally isolated from the red flour beetle. *Tc Buster* has been optimized for transposition in mammalian cells and a hyperactive mutant (TcB-M) has been evolved by Bio-Techne (Minneapolis, MN). We obtained wild-type TcB and TcB-M mRNA from Bio-Techne and used them to deliver a model expression cassette containing a second generation CD19-CAR, mutant DHFR, and EGFP flanked by *Tc Buster* inverted terminal repeats (ITRs) to primary human PB NK and T cells.

The use of transposons in NK cells has been limited by DNA toxicity^1^. To avoid this, we delivered the transposase as modified mRNA^**57**^ and the transposon via nanoplasmid vectors which have a small backbone, high supercoiling, and are readily scaled under cGMP compliance^**58**^. We also optimized activation, electroporation, recovery, and expansion conditions to achieve 49.63% (± 4.64%) integration efficiency in NK cells without selection (Figure 1). One such modification we made to our protocol included pre-treating NK cells with an RNase inhibitor prior to electroporation. This enhanced transposition efficiency from 24.05% (±1.36%) to 49.63% (±4.64%). This is likely due to degradation of the transposase mRNA by RNases produced by NK cells without RNase inhibition. However, there is very limited data available on the expression of ribonucleases in NK cells in resting or activated states. Because of this, we used a broad RNase inhibitor which targeted RNase A, RNase B, and RNase T2 in our workflow^**51**^. In future studies, it would be useful to profile NK cells for expression of a full panel of RNases at resting state and during several activation timepoints, as there may be other RNases produced by NK cells that could be targeted to further enhance efficiency.

We included a MTX-resistant DHFR mutein (L22F, F31S) in our cargo^**54**^. This allowed us to enrich our engineered population using MTX, without the need for GMP-compliant cell sorting facilities or clinical-grade monoclonal antibodies. Further, MTX is approved for clinical use and is being as described here for ex vivo enrichment of engineered cells (NCT04483778) (Figure 2). Overall, we developed a CAR-NK cell manufacturing protocol that includes 4 days of feeder cell activation before electroporation, re-expansion 2 days after electroporation, and a third expansion with MTX selection (Figure 2A). This process results in ∼1400-fold expansion from electroporation input when cells are engineered with TcB-M (Figure 2D). Current clinical trials using CAR-NK cells have treated patients with doses ranging from 1×10^5^ – 1×10^7^ cells/kg of body weight^**59**^. Thus, the average North American patient (80 kg) would receive a dose of 8×10^6^ NK cells on the low end and 8×10^8^ NK cells on the high end. A single dose of 1×10^7^ cells/kg could be produced in 20 days from a starting number of just 5.8×10^5^ NK cells. We routinely obtain apheresis products from healthy donors containing an average of 1×10^9^ NK cells (n=12 donors). Thus, we could generate an average of 1724 high (1×10^7^ cells/kg) doses from each donor.

Importantly, we validated the function of CAR-NK cells produced using our protocol in standard *in vitro* assays. CAR-NK cells showed high levels of degranulation, inflammatory cytokine production, and target cell killing compared to CAR-negative controls (Figure 3A – 3D). Interestingly, it has been shown that cryopreservation may lead to reduced cytotoxicity of NK cells^**55**^. An off-the-shelf product requires cryopreservation and banking of engineered cells. We did observe reduced cytotoxicity of CAR-NK cells immediately after thaw, but this phenotype was rescued after an overnight rest in media containing 100 IU/mL IL2 (Figure 3E). However, it is likely not feasible to include this type of overnight rest period in a clinical protocol. We observed efficient target cell killing immediately after thaw when we used higher E:T ratios. Therefore, instead of implementing an overnight rest in a clinical protocol, it is likely more practical to simply deliver a higher dose of cells. This is feasible as the administration of CAR-NK cells clinically has not been associated with cytokine release syndrome, neurotoxicity, or graft-versus-host disease (GvHD), and the maximum tolerated dose has not been reached^**59**^.

Finally, we deployed the Tc Buster system for the production of CAR-T cells. CAR-T cells have been generated for preclinical and clinical use with a number of transposon systems including SB and PB^**25–28,30–32,35,36,43**^. Here, we demonstrated the feasibility and high efficiency of their production using *Tc Buster* and validated their subsequent effector function (Figure 4).

Our work provides a platform for robust delivery of multicistronic, large cargo via transposition to primary human PB NK and T cells. Our results demonstrate that CAR-expressing NK and T cells can be enriched using MTX selection at clinically relevant doses, while maintaining high viability and function, and we can manufacture a large number of doses in a matter of three weeks. This approach represents a versatile, safe, and cost-effective option for the manufacture of CAR-NK and CAR-T cells compared to viral production methods. Importantly, further studies are required to test Tc Buster delivery of larger cargo that exceeds the carrying capacity of viral vectors. Future work will focus on the validation of CAR-NK and CAR-T cells in preclinical *in vivo* models. We will also explore the use of other CARs targeting solid tumors and combining CAR-NK and CAR-T cells in a single infusion.

## Materials and Methods

### Vectors and Reagents

*Tc Buster* (TcB) and Hyperactive *Tc Buster* (TcB-M) sequences were obtained from Bio-Techne (Minneapolis, MN). CD19-DHFR-EGFP flanked by *Tc Buster* ITRs was cloned into a Nanoplasmid backbone (Nature Technology, Lincoln, NE).

### Donor T and NK Cell Isolation

Peripheral blood mononuclear cells (PBMCs) from de-identified healthy human donors were obtained by automated leukapheresis (Memorial Blood Centers, Minneapolis, MN). CD3+ T cells or CD56+CD3-NK cells were isolated from the PBMC population using the EasySep Human T Cell Isolation Kit or EasySep Human NK Cell Isolation Kit (STEMCELL Technologies, Cambridge, MA). T cells were frozen at 1-2 × 10^7^ cells/mL and NK cells were frozen at 5 × 10^6^ cells/mL in CryoStor CS10 (STEMCELL Technologies, Cambridge, MA) and thawed into culture as needed. Samples were obtained after informed consent with approval from the University of Minnesota Institutional Review Board (IRB 1602E84302)

### T Cell Culture

T cells were cultured in OpTimizer CTS T Cell Expansion SFM containing 5% CTS Immune Cell SR (ThermoFisher, Waltham, MA), L-Glutamine, Penicillin/Streptomycin (Lonza, Basel, Switzerland), 10 mM N-Acetyl-L-cysteine (Sigma-Aldrich, St. Louis, MO), 300 IU/mL IL-2, 5 ng/mL IL7, and 5 ng/mL IL-15 (PeproTech, Rocky Hill, NJ). T cells were activated with DynaBeads Human T-Activator CD3/CD28 (ThermoFisher, Waltham, MA) at a 2:1 bead:cell ratio for 48 hours prior to electroporation. Following electroporation, T cells were re-stimulated with DynaBeads and maintained at ∼1 × 10^6^ cells/mL.

### NK Cell Culture

NK cells were cultured in CTS AIM V SFM containing 5% CTS Immune cell SR (ThermoFisher, Waltham, MA), Penicillin/Streptomycin, and IL-2 (100 IU/mL). NK cells were activated by co-culture with X-irradiated (100 Gray) feeder cells (K562 expressing membrane-bound IL21 and 41BB-L^**60**^) at indicated feeder:NK ratios (2:1 prior to electroporation, 5:1 48 hours after electroporation, or 1:1 for all subsequent expansions).

### T Cell Electroporation

After 48 hours of stimulation, DynaBeads were magnetically removed, and T cells were washed once with PBS prior to resuspension in electroporation buffer. The 4D-Nucleofector (Lonza, Basel, Switzerland) and P3 kit was used with 1 × 10^6^ T cells per 20 µL cuvette, 1 µg transposase mRNA, 1 µg transposon nanoplasmid, and the Nucleofector program FI-115. Transposon nanoplasmid alone was used as a control for all experiments. T cells were allowed to recover in antibiotic-free medium containing 1 µg/mL DNase I solution (STEMCELL Technologies, Cambridge, MA) at 37 °C, 5% CO2 for 30 minutes following gene transfer, and then were cultured in complete T cell medium and re-stimulated with DynaBeads.

### Electroporation of activated NK cells

Feeder cell-activated NK cells were washed once with PBS and resuspended at 3 × 10^7^ cells/mL in electroporation buffer. Protector RNase inhibitor (Sigma Aldrich, St. Louis, MO) was added to the mixture at a concentration of 0.8 U/µL and incubated for 5 minutes at room temperature. The cell mixture was added to 1 µg of transposase mRNA and 1 µg transposon nanoplasmid on ice. Transposon nanoplasmid alone was used as a control for all experiments. This mixture was electroporated in a 10 µL tip using the Neon Transfection System (ThermoFisher, Waltham, MA) under the following conditions: 1850 volts, pulse width of 10 ms, two pulses. NK cells were allowed to recover at a density of 1.5 × 10^6^ cells/mL in antibiotic-free medium containing 1 ug/mL DNase I solution (STEMCELL Technologies, Cambridge, MA), and were then cultured in complete NK cell medium at a density of 6 × 10^5^ cells/mL. 48 hours after electroporation, NK cells were expanded with feeder cells at a 5:1 feeder:NK ratio.

### Antibodies and Flow Cytometry

The following antibodies, proteins, and dyes were used: APC- or PE-conjugated anti-CD56 (clone REA196; Miltenyi Biotec), PE-conjugated anti-CD3 (clone SK7; BD Biosciences), Atto 647N-conjugated recombinant human CD19 (BioTechne), Brilliant violet 421-conjugated anti-IFNγ (clone 4S.B3; BioLegend), APC-labeled anti-TNFa (clone Mab11, BioLegend), Brilliant violet 650-conjugated anti-CD107a (clone H4A3; BD Biosciences), SYTOX Blue dead cell stain (ThermoFisher), Fixable viability dye eFluor 780 (eBioscience). Flow cytometry assays were performed on a CytoFLEX S flow cytometer (Beckman Coulter) and all data were analyzed with FlowJo verson 10.4 software (FlowJo LLC).

### NK Cell Functional Assays

For intracellular cytokine staining, NK cells were plated at 2.5 × 10^6^ cells/mL in NK cell medium without cytokines. After incubation overnight, the CD19+ Burkitt’s Lymphoma cell line Raji was added at the indicated effector-to-target (E:T) ratios. Brilliant violet Anti-CD107a was added to the culture and cells were incubated for 1 hour at 37 °C. Brefeldin A and monensin (BD Biosciences, San Jose, CA) were added and cells were incubated for an additional 4 hours. Cells were stained with fixable viability dye, then for extracellular antigens. Cells were fixed and permeabilized using BD Cytofix/Cytoperm (BD Biosciences, San Jose, CA) following manufacturer’s instructions. Cells were then stained for intracellular IFNγ and TNFα and analyzed by flow cytometry.

### Target Cell Killing Assays

T cells or NK cells were cultured overnight in medium without cytokines. Luciferase-expressing Raji cells were seeded into a black round-bottom 96-well plate (3 × 10^4^ cells per well). T cells or NK cells were added to the wells in quadruplicate at the indicated E:T ratios. Target cells without effectors served as a negative control (spontaneous cell death) and target cells incubated with 1% NP-40 served as a positive control (maximum killing). Co-cultures were incubated at 37C for 24 hours. After incubation, D-luciferin (potassium salt; Gold Biotechnology, St. Louis, MO) was added to each well at a final concentration of 25 ug/mL and incubated for 10 minutes. Luminescence was read in endpoint mode using a BioTek Synergy microplate reader.

### Statistical Analysis

The Student’s t-test was used to test for significant differences between two groups. Differences between 3 or more groups were tested by one-way ANOVA analysis followed by Tukey’s *post-hoc* test. All assays were repeated in 3-5 independent donors. Mean values +/- standard error of the mean (SEM) are shown. The level of significance was set at α = 0.05. Statistical analyses were performed using GraphPad Prism 8.0.

## ACKNOWLEDGEMENTS

We thank Bio-Techne for providing us with TcB-M and their support of these studies.

## AUTHOR CONTRIBUTIONS

E.J.P, B.R.W, and B.S.M conceived of the project, designed experiments, and directed the research. E.J.P performed all experiments with assistance from W.S.L, B.J.W, N.J.S, and J.G.S. B.J.W and N.J.S performed T cell culture and electroporations. E.J.P performed all data analysis. E.J.P wrote the manuscript with input from all authors.

## Supplemental Material

**Supplemental Figure 1.**
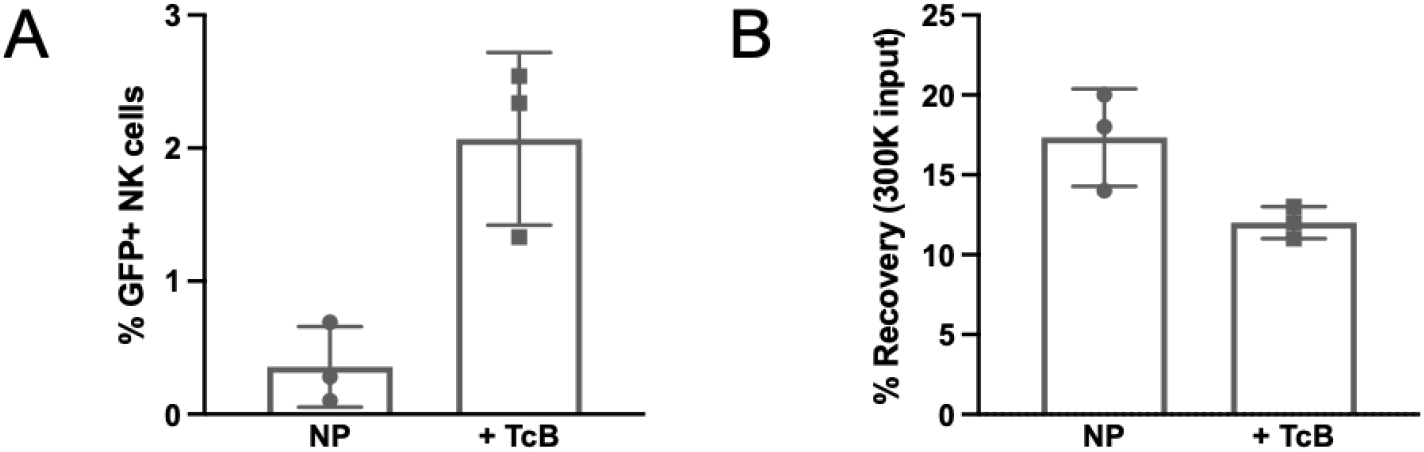
Transposition efficiency and cell health are poor when cells are electroporated on day 7 of activation. Primary human peripheral blood (PB) NK cells were expanded for 7 days with mbIL21- and 41BBL-expressing K562 feeder cells at a 2:1 (feeder:NK) ratio. NK cells were electroporated with the nanoplasmid transposon (NP) alone or in combination with mRNA encoding *Tc Buster* (TcB). Two days after electroporation, NK cells were expanded with feeder cells (1:1 feeder:NK ratio) for 1 week to allow for the loss of transient NP expression. After this expansion, GFP expression was measured by flow cytometry (A) and cells were counted to calculate recovery from electroporation input (B).

**Supplemental Figure 2.**
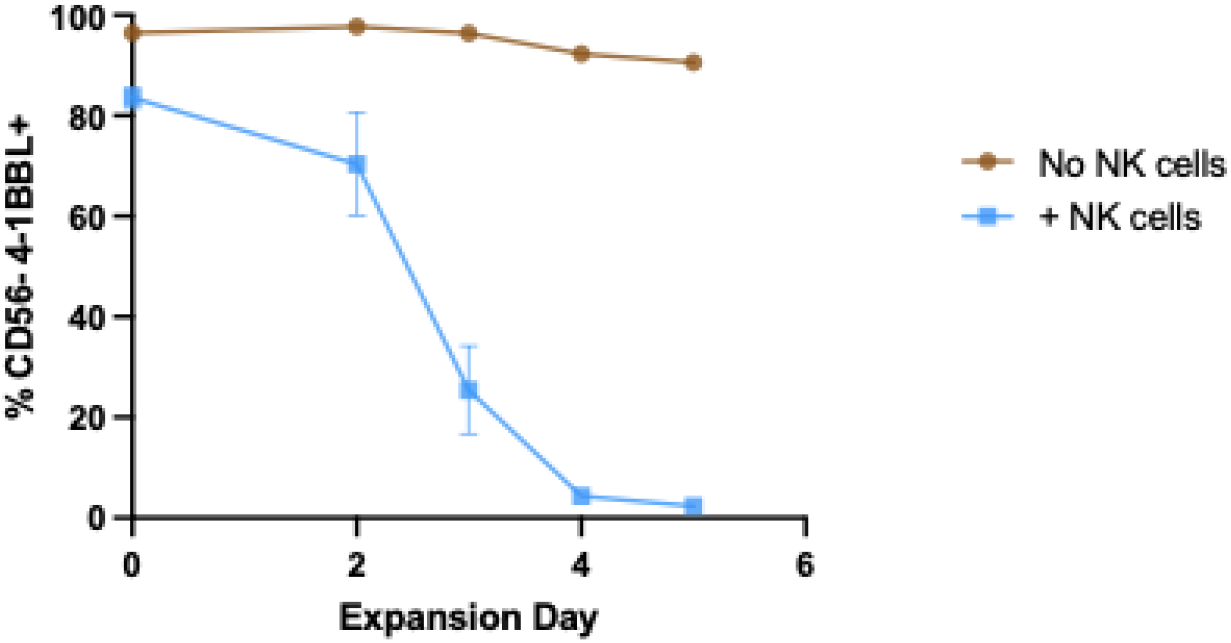
Feeder cells are eliminated during co-culture with NK cells. K562 feeder cells expressing mbIL21 and 41BBL were irradiated (100 Gray) and cultured alone or in the presence of primary human peripheral blood NK cells (feeder-to-NK ratio = 5:1). Cultures were stained with antibodies for 41BB-L and NK cell marker CD56 and analyzed by flow cytometry. N=3 human NK cell donors.

**Supplemental Figure 3.**
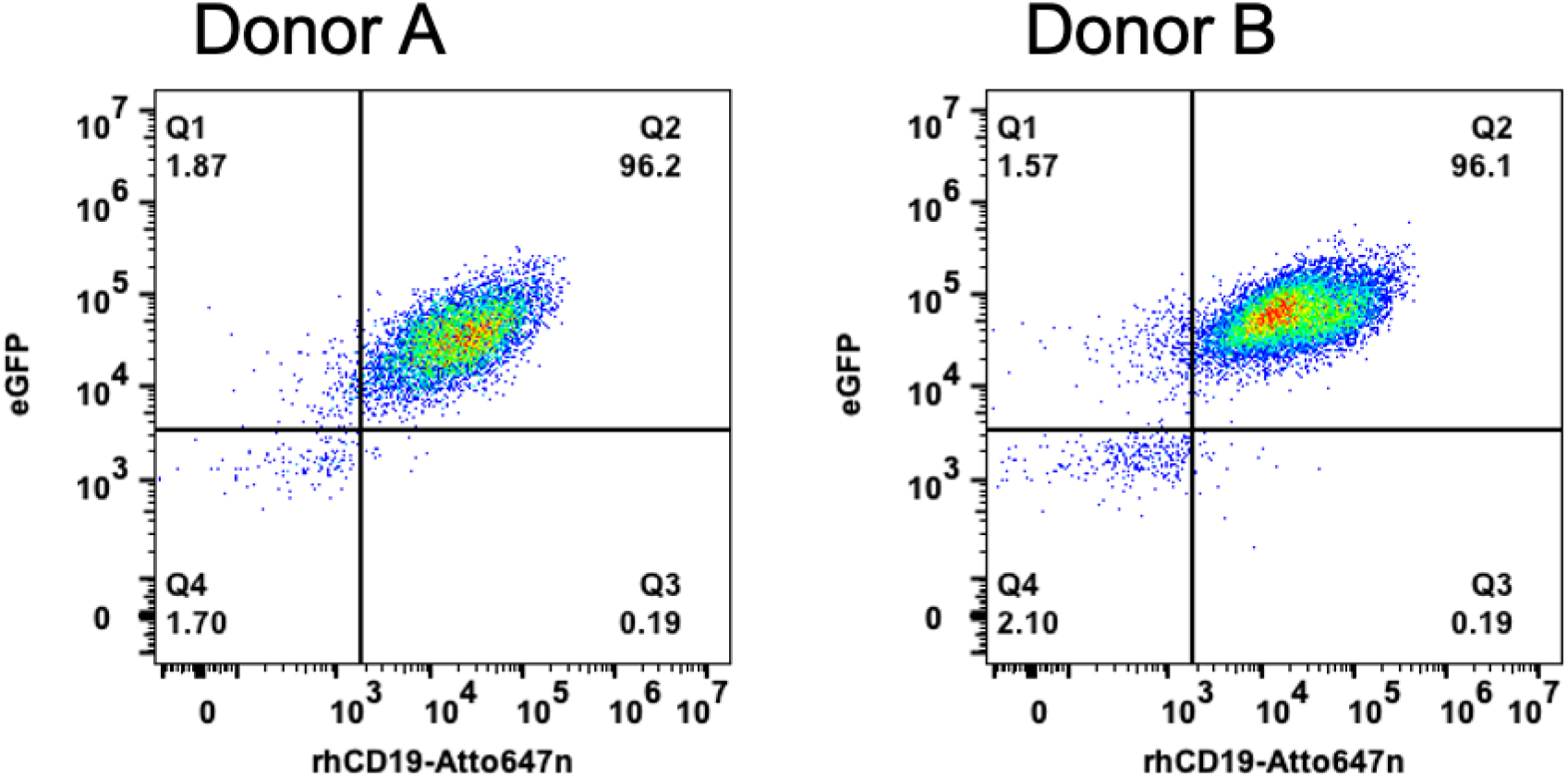
Representative flow plots showing CD19-CAR staining using recombinant human CD19 (rhCD19) conjugated to fluorescent Atto 647N. CAR-NK cells were engineered usingTcB-M and selected with 250 nM MTX. Cells were incubated with rhCD19 Atto 647N and analyzed by flow cytometry.

**Supplemental Figure 4.**
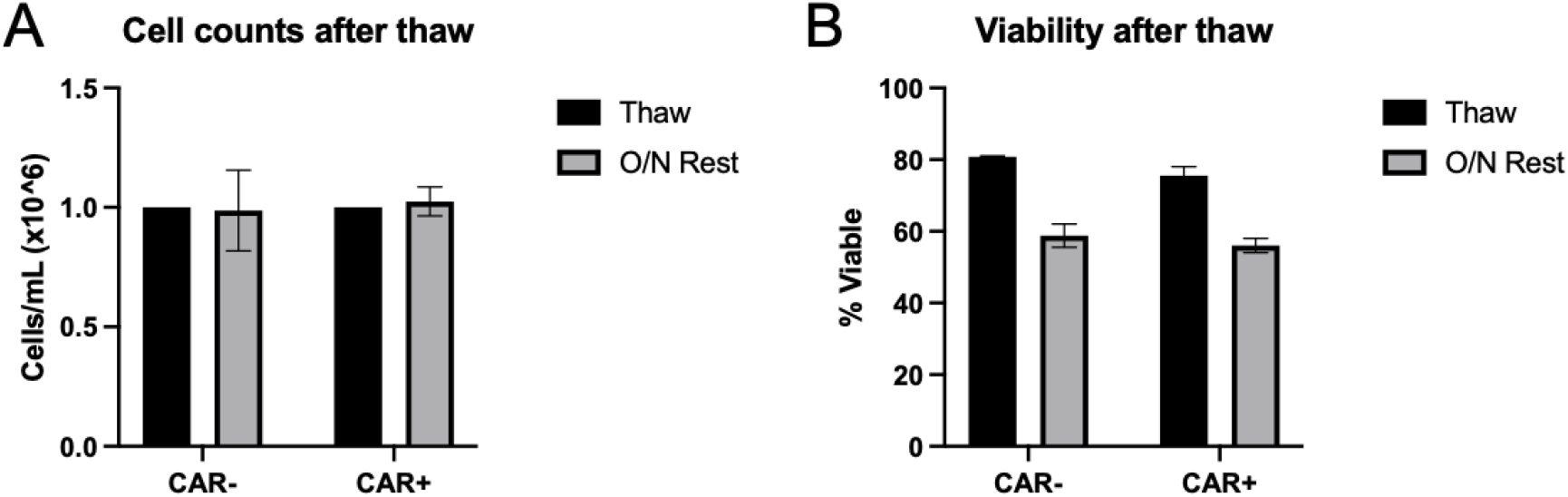
NK cell viability and recovery after cryopreservation. CAR-positive and CAR-negative NK cells were cryopreserved after the 20-day production timeline. After thaw, cells were counted (A), and viability was measured by trypan blue exclusion (B). Cells were immediately plated in killing assays or plated at a density of 1×10^6^ cells/mL in media containing 100 IU/mL IL2 overnight. After overnight rest, cells were counted, and viability was measured again.

